# Single-cell chromatin accessibility in glioblastoma delineates cancer stem cell heterogeneity predictive of survival

**DOI:** 10.1101/370726

**Authors:** P. Guilhamon, M.M. Kushida, A. Nikolic, D. Singhal, G. MacLeod, S.A. Madani Tonekaboni, F.M.G. Cavalli, C. Arlidge, N. Rajakulendran, N. Rastegar, X. Hao, R. Hassam, L.J. Smith, H. Whetstone, F.J. Coutinho, B. Nadorp, K.I. Ellestad, H.A. Luchman, J.A. Chan, M.S. Shoichet, M.D. Taylor, B. Haibe-Kains, S. Weiss, S. Angers, M. Gallo, P.B. Dirks, M. Lupien

## Abstract

Chromatin accessibility discriminates stem from mature cell populations, enabling the identification of primitive stem-like cells in primary tumors, such as Glioblastoma (GBM) where self-renewing cells driving cancer progression and recurrence are prime targets for therapeutic intervention. We show, using single-cell chromatin accessibility, that primary GBMs harbor a heterogeneous self-renewing population whose diversity is captured in patient-derived glioblastoma stem cells (GSCs). In depth characterization of chromatin accessibility in GSCs identifies three GSC states: Reactive, Constructive, and Invasive, each governed by uniquely essential transcription factors and present within GBMs in varying proportions. Orthotopic xenografts reveal that GSC states associate with survival, and identify an invasive GSC signature predictive of low patient survival. Our chromatin-driven characterization of GSC states improves prognostic precision and identifies dependencies to guide combination therapies.

## INTRODUCTION

Glioblastoma (GBM) is a lethal form of brain cancer with standard surgery and radiation giving a median survival of only 12.6 months[1]. The addition of temozolomide chemotherapy provides only an additional 2.5 months in the small subset of responsive patients[2]. Despite extensive characterization and stratification of the bulk primary tumors, no targeted therapies have been successfully developed[1,3]. GBM tumors are rooted in self-renewing tumor-initiating cells commonly referred to as glioblastoma stem cells (GSCs)[4] that drive disease progression *in vivo[5,6]* and display resistance to chemo- and radiotherapy leading to disease recurrence[7]. The promise of therapeutically targeting self-renewing tumor-initiating cancer cells depends on our capacity to capture the full range of heterogeneity within this population from individual tumors. Intratumoral heterogeneity within primary GBM has recently been documented through single cell RNA-seq experiments and revealed a continuum between four cellular states[8]: neural-progenitor-like (NPC), oligodendrocyte-progenitor-like (OPC), astrocyte-like (AC), and mesenchymal-like (MES)[8]. A subsequent study[9] using single-cell gene-centric enrichment analysis placed GBM cells along a single axis of variation from proneural to mesenchymal transcriptional profiles, with cells expressing stem-associated genes lying at the extremes of this axis. Hence, primary GBM consists of distinct states, across which stem-like cells appear to be found. Whether these stem-like cells found across GBM states represent functionally distinct GSC populations with tumor-initiating properties and unique dependencies remains to be established to guide therapeutic progress. To address this issue, we combined single-cell technologies to define GSC composition in primary GBM with functional assays to reveal the unique dependencies across GSCs, reflective of invasive, constructive and reactive states that relate to patient outcome.

## RESULTS

Chromatin accessibility readily discriminates stem from mature cell populations[10], which can be resolved at the single cell level through single-cell ATAC-seq[11,12] taking into account non-gene centric features, such as accessibility of noncoding elements and total amount of accessible DNA sequences. Applying single-cell chromatin accessibility profiling (scATAC-seq) across four *IDH* wild-type primary adult GBM tumors (3797 cells) revealed 7-9 accessibility modules in each tumor based on unsupervised clustering (Fig. 1A). We assigned cells to each of the four scRNA-seq-derived cellular states[8] based on individual cells’ chromatin accessibility enrichment scores for the promoter regions of each state’s signature genes. The MES state reported from scRNA-seq[8] dominates the identity of two or more modules reported from chromatin accessibility in every tumor (Fig. 1B-C). In contrast, the NPC and OPC states are mixed within the same module defined based on chromatin accessibility, dominating over the other states typically in at least two modules. Cells assigned to the AC state did not preferentially cluster within a single module reported from chromatin accessibility (Fig. 1B-C). Collectively, our results suggest that chromatin accessibility reflects a greater stratification of the MES state, detects similarities between the OPC and NPC states and heterogeneity within the AC state.

**Fig. 1.**
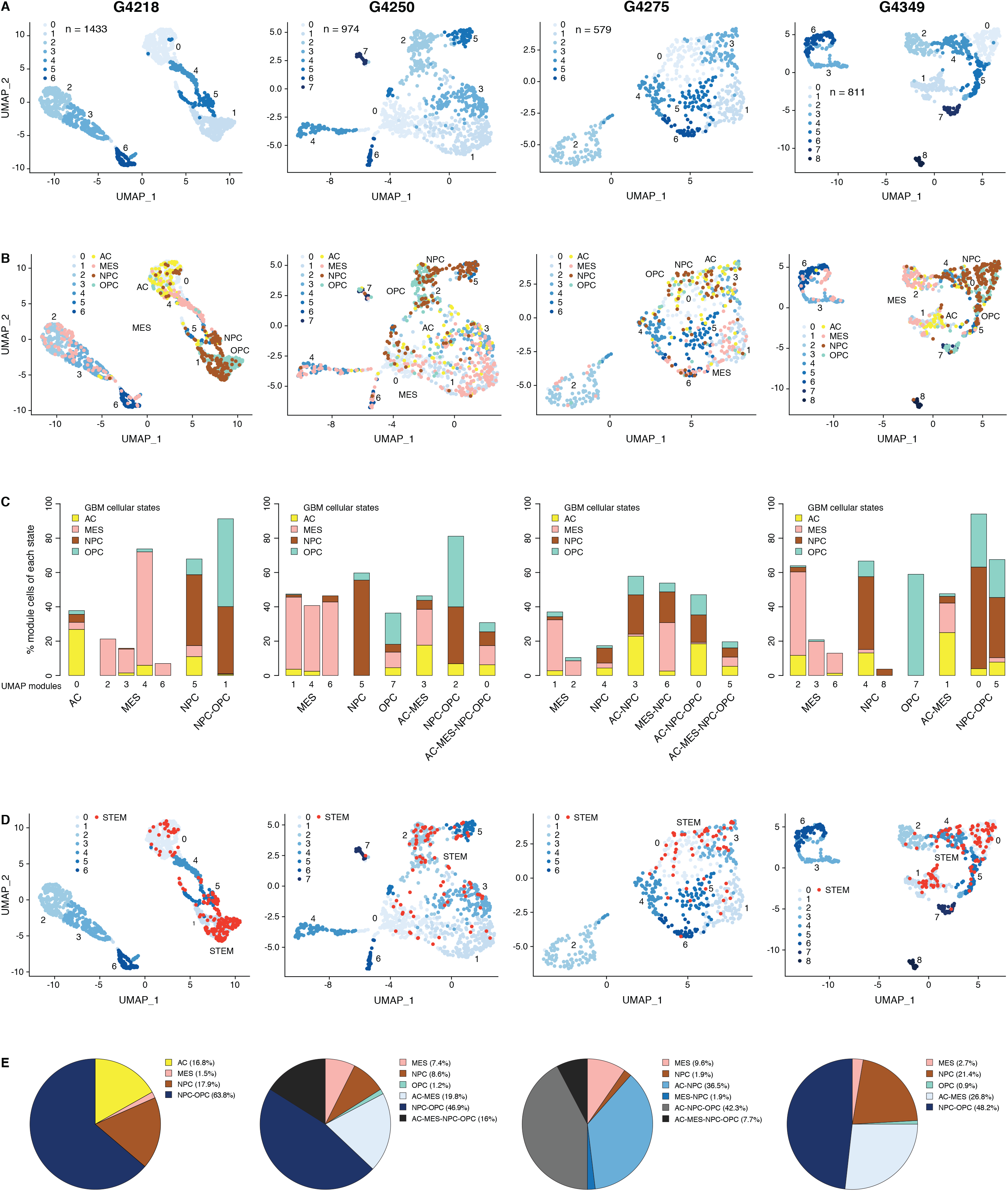
The diverse GBM cancer stem cell pool. **(A)** UMAP representation of chromatin accessibility across four primary GBM. **(B)** UMAPs with tumor cells assigned to cellular states. **(C)** UMAP modules are grouped by dominant cellular state. **(D)** UMAPs with cancer stem cells highlighted based on the enrichment of GBM cancer stem transcription factor promoters. **(E)** Distribution of cancer stem cells across the modules dominated by each cellular state.

To identify putative cancer stem cells within each primary tumor, we next focused on the level of chromatin accessibility at promoters of 19 transcription factors previously associated with self-renewal and tumor-propagating capacity in GBM[13] (Fig. 1D). Individual cells scoring as putative cancer stem cells were not restricted to a unique module defined by chromatin accessibility but were distributed across a subset of modules, suggesting heterogeneity across cancer stem cells in primary GBM, in agreement with reports relying on single GBM cell labelling[6,10,12–20] assessing the heterogeneity of self-renewing tumor-initiating cells. Putative cancer stem cells identified in primary GBM through scATAC-seq were found in modules ascribed to every one of the four cellular states defined by gene expression[8], predominantly within NPC and OPC containing modules and a smaller fraction (<10%) in MES-specific modules across all four tumors (Fig. 1E). This suggests that the core transcriptional unit of cancer stem cells in primary GBM[13] is not restricted to a unique population defined by its global transcriptional or chromatin accessibility profile with the resolution achieved with current single-cell technologies.

To further probe the heterogeneity in chromatin accessibility within the GBM cancer stem cell pool, we derived GSC populations from 27 adult *IDH* wild-type GBM tumors[21] and profiled their chromatin accessibility by bulk ATAC-seq (Fig. 2A). Each patient-derived GSC showed a similar enrichment for accessible chromatin regions in promoters and 5’UTRs, and depletion in introns and distal intergenic regions (Fig. 2B). Collectively, we uncovered 92% of the total predicted regions of accessible chromatin (255,891 regions) within GSCs based on a saturation analysis using a self-starting nonlinear regression model across the 27 samples (Fig. 2C). We next assessed the similarity between these GSCs and the putative cancer stem cells found by scATAC-seq in the four primary GBMs. GSCs were identified within each tumor by calculating the enrichment of accessible chromatin regions shared by a majority of GSCs (>14/27) in each tumor cell (Fig. 2D). Comparing the distribution of GSCs across the 7-9 modules defined by scATAC-seq to that of the 19 transcription factor-derived cancer stem cell signature demonstrates concordance between the two signatures (Fig. 2E). Moreover, the enrichment z-scores for both cancer stem signatures (ie stem transcription factors signature and GSC chromatin accessibility signature) are significantly correlated across cells in all four tumors (p ≤ 1.6e^−5^) (Fig. 2F). Additionally, an average of 91.2% (85.1-100%) of the cells identified by either signature display the hallmark GBM copy number changes at chromosomes 7 and 10, confirming their neoplastic status (Fig. S1A-D). Collectively, these results demonstrate that the patient-derived GSC populations reflect the chromatin identity of putative cancer stem cells found in primary brain tumors, highlighting the value of these GSCs as models to deepen our understanding of individual cells within primary GBM with features found in self-renewing tumor-initiating cells. Accordingly, spectral clustering of the 27 patient-derived GSC ATAC profiles identifies three distinct states of self-renewing tumor-initiating cells (Fig. 3A). Expression profiling of these 27 GSCs by RNA-seq reveals GSCs significantly enriched for the signatures of each of the three TCGA GBM subtypes (proneural, classical, and mesenchymal)[22] (Fig. 3B, top panel). However, the assignment of the proneural, classical and mesenchymal subtypes across GSCs did not match the three clusters identified from ATAC-seq (Fig. 3B, top panel). Conversely, clustering the GSCs by gene expression, independently of their chromatin accessibility, did largely recapitulate the GSC states defined by chromatin accessibility (Fig. 3B, bottom panel). This suggests that the mismatch between the GSC cluster from chromatin accessibility profiles and TCGA expression subtypes is not mainly due to differences between chromatin accessibility and gene expression. A more likely possibility is that given the TCGA subtypes were determined from bulk GBM, they may not fully capture the nature of rarer populations found within a tumor, such as the cancer stem cell populations.

**Fig. 2.**
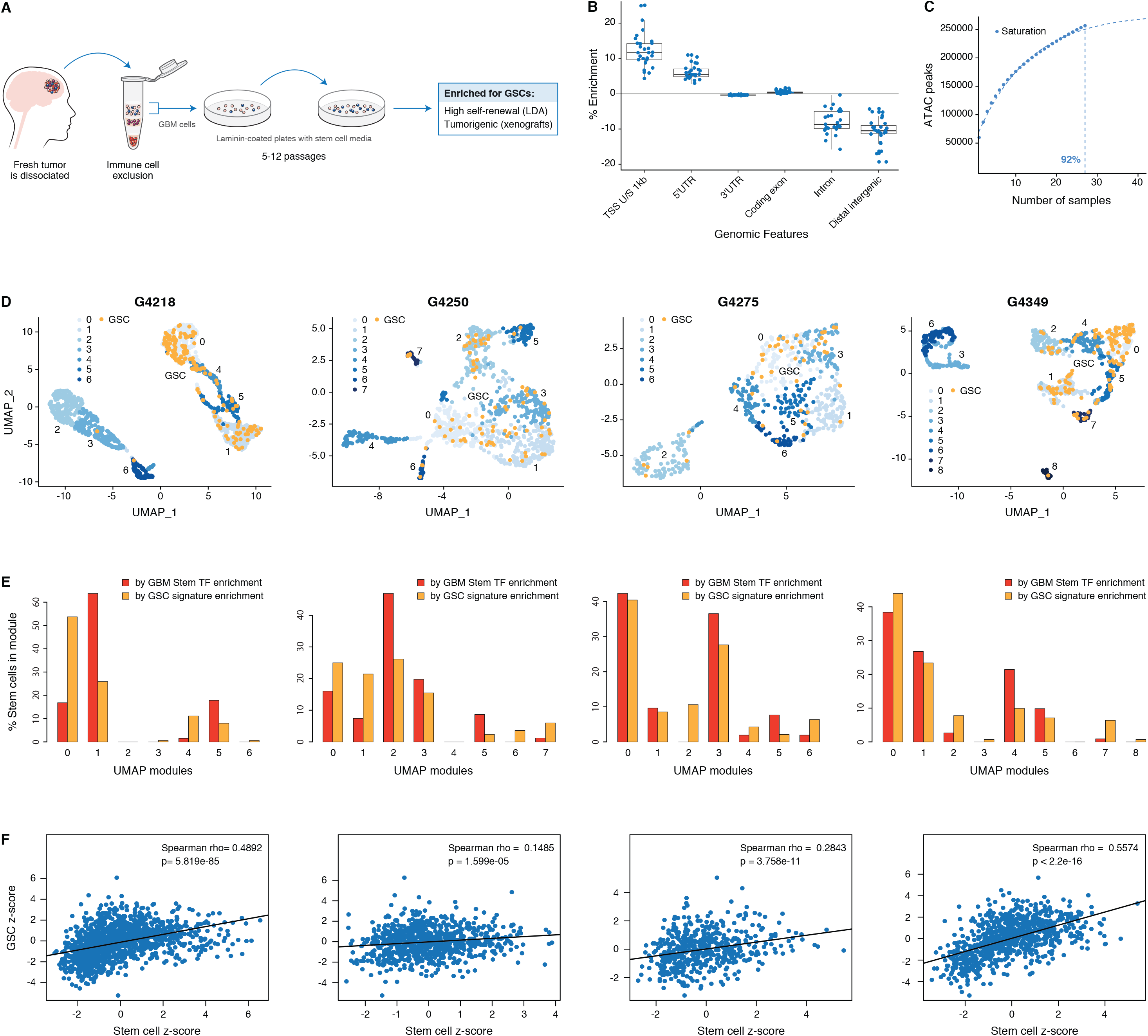
GSCs recapitulate the GBM cancer stem cell population. **(A)** Schematic representation of the GSC derivation process, from patient tumor to GSC-enriched population. **(B)** Genomic feature enrichment of accessible chromatin peaks. **(C)** Saturation curve for the 27 GSCs. **(D)** UMAPs with GSCs highlighted based on the enrichment of shared accessible regions across GSCs. **(E)** Proportion of UMAP modules assigned to cancer stem cells and GSCs. **(F)** Correlation of z-scores for each signature for each cell in each primary GBM.

**Fig. 3.**
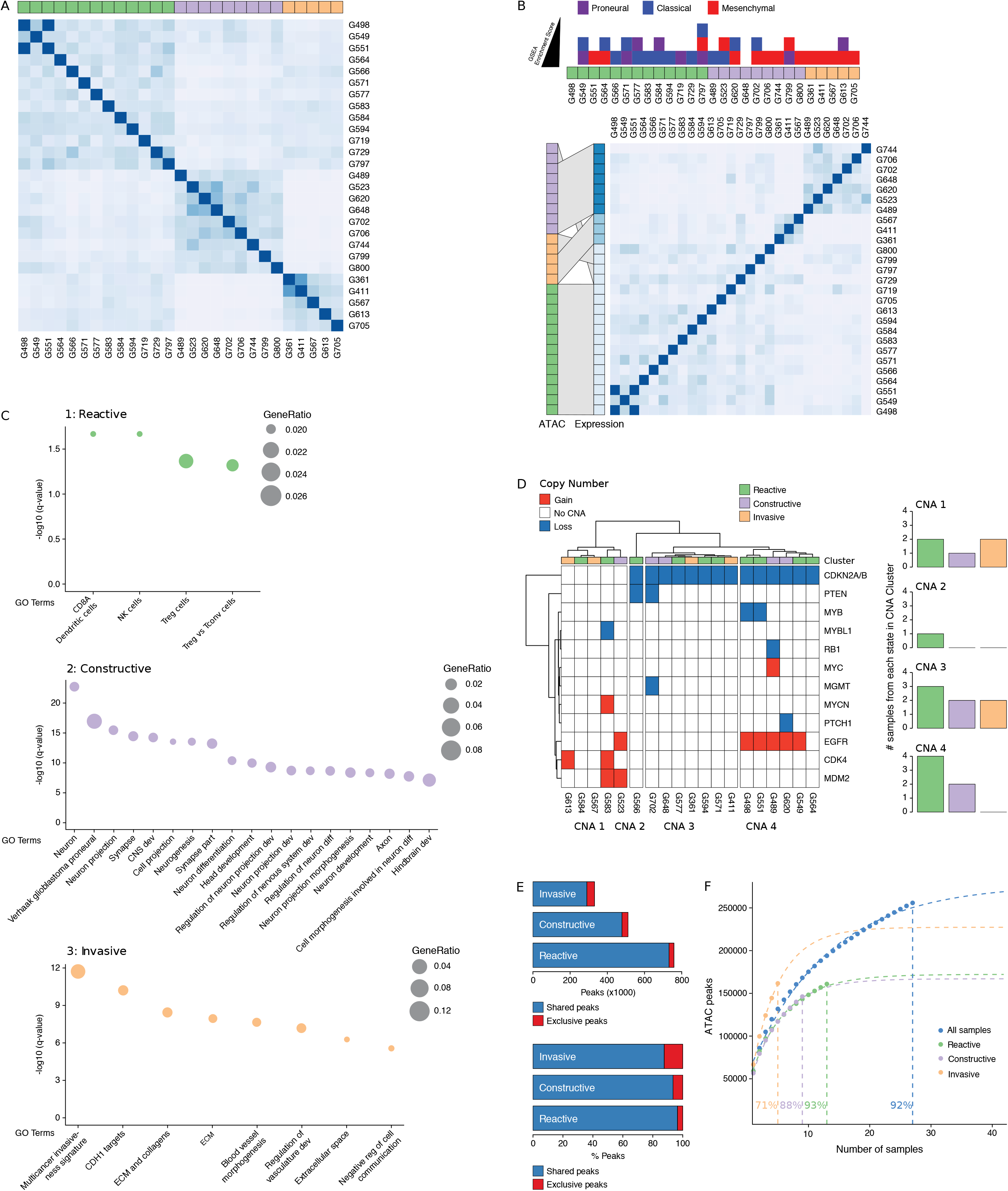
Three GSC states driven by chromatin accessibility. **(A)** Spectral clustering of ATAC-seq signal across peaks in 27 GSCs. **(B)** Top: enrichment of TCGA subtypes across GSCs and comparison to GSC states; Bottom: spectral clustering of gene expression across GSCs and comparison to chromatin-derived GSC states. **(C)** Gene set enrichment analysis in each GSC state. **(D)** CNAs across GSCs identified from DNA methylation array data cluster GSCs into four subgroups. **(E)** Number and percentage of peaks unique and shared in each GSC state. **(F)** Saturation analysis of each individual state.

Gene set enrichment analysis with GSEA[23] and g:profiler[24] using genes exclusively enriched for both expression and promoter chromatin accessibility in each subtype reported significantly enriched terms defining the largest GSC state as a Reactive state, with terms related to immune cells and response (Fig. 3C, top panel). A second GSC state was enriched for Constructive gene sets involved in brain, neuron, and glial cell development (Fig. 3C, middle panel). The third and smallest GSC state presented an Invasive state characterized by terms relating to the extracellular matrix and angiogenesis (Fig. 3C, bottom panel). We next mapped copy number alterations (CNAs) across the 27 GSCs by applying the molecular neuropathology classifier tool[25] to DNA methylation data from the GSCs (Fig. 3D). While common GBM CNAs, including EGFR gains and CDKN2A/B loss, were observed, the CNA-based classification of GSCs failed to match the three chromatin accessibility-derived states, suggesting the three GSC states are not genetically defined (Fig. 3D). Further comparison of the accessible chromatin in each GSC state reveals that only a small subset of accessible chromatin regions drives the three GSC states (Fig. 3E and Fig. S1E-G). Our ability to discriminate GSC state-specific regions of accessible chromatin is reflective of the comprehensiveness of our cohort to saturate the detection of accessible regions to 93%, 88%, and 71% across the Reactive (n=13), Constructive (n=9), and Invasive (n=5) state GSCs, respectively (Fig. 3F).

Considering that regions of accessible chromatin serve as binding sites for transcription factors engaging in gene expression regulation, we next tested for DNA recognition motif family enrichment across regions exclusively accessible in Reactive, Constructive or Invasive GSC states (Fig. 4A and Fig. S2A). The most enriched DNA recognition motif families in each state were either depleted or showed low-level enrichment in the other states. Specifically, the DNA recognition motifs for the interferon-regulatory factor (IRF) and Cys2-His2 zinc finger (C2H2 ZF) transcription factor families were enriched in the Reactive state (Fig. 4A, top panel). Regulatory factor X (RFX) and basic helix-loop-helix (bHLH) DNA recognition motifs were enriched in the Constructive state (Fig. 4A, middle panel), while the Forkhead motif family was enriched in the Invasive state (Fig. 4A, bottom panel). Genome-wide CRISPR/Cas9 essentiality screens (Fig. S2B) in three Reactive, two Constructive and one Invasive GSC[26] revealed the preferential requirement for expressed transcription factors (Fig. S2C-E) recognizing the enriched DNA recognition motif in a state-specific manner (Fig. 4B). Specifically, the SP1 regulatory network is preferentially essential in the Reactive state GSCs (Fig 4B, top panel), ASCL1, OLIG2, AHR, and NPAS3 are uniquely essential in the Constructive state GSCs (Fig. 4B, middle panel) and FOXD1 in essential only in the Reactive state GSC (Fig. 4B, bottom panel). Notably, SP1 itself is exclusively essential in only one Reactive GSC (G564). However, of the 36 transcription factors from the Reactive-enriched families (IRF and C2H2 ZF) that were essential in at least one Reactive GSC and not in any of the Constructive or Invasive GSCs, 13 are directly regulated by SP1 (Fig. S2C), thus suggesting that the SP1 regulatory network as a whole, rather than SP1 on its own, is key in the Reactive GSC state. Notably, all six transcription factors display significantly higher expression in GBM compared to normal brain[27] (Fig. S2F), further supporting their function as key regulators of tumor initiation and development. An additional gene set enrichment analysis combining genes exclusively essential to each state with the putative targets of the key transcription factors outlined above identifies additional enriched terms supporting the identities of the three GSC states as Reactive, Constructive, and Invasive (Fig. S2G).

**Fig. 4.**
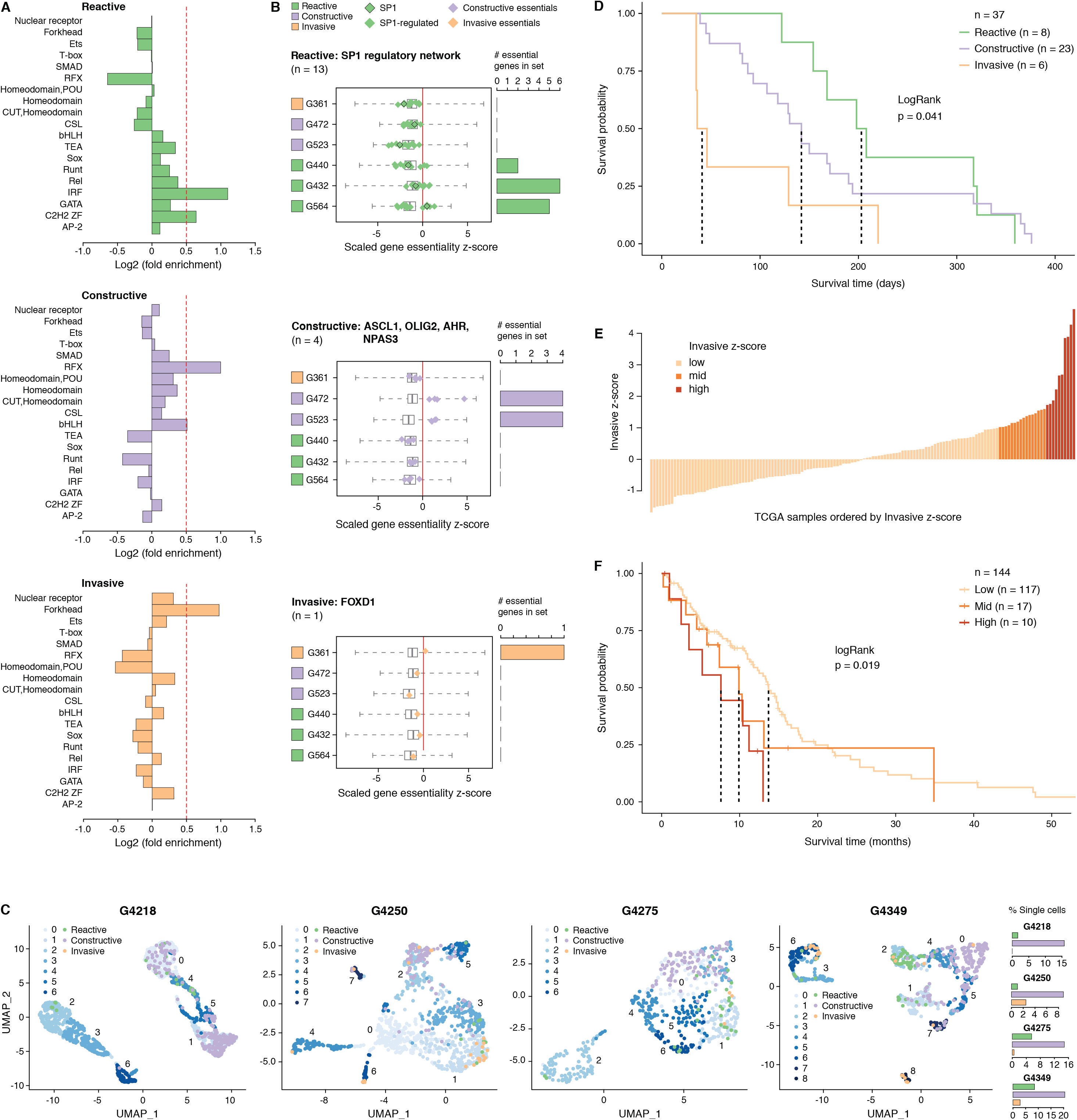
Functional diversity between GSC states drives survival in GBM. **(A)** Motif family enrichment in each cluster; log2(Fold Enrichment) > 0.5 threshold selected based on the distribution of values in each cluster (Fig. S1A). **(B)** Z-score distribution of key essential genes in each cluster. Red line corresponds to the empirically determined threshold for essentiality in each tested line, scaled and adjusted to zero. Boxplot whiskers in this case extend to data extremes. Side barplots show the total count of the key cluster-specific regulators found essential in each subtype. **(C)** UMAPs with GSCs from each state highlighted based on the enrichment of the top differentially accessible regions in each GSC state. **(D)** Kaplan-Meier plot for orthotopic xenografts grouped by GSC state. The dotted lines indicate median survival. The pairwise p-values are also significant for Invasive vs Reactive (p=0.02) and Invasive vs Constructive (p=0.045) but not for Reactive vs Constructive (p=0.45). **(E)** TCGA samples ordered by increasing concordance with Invasive GSCs and grouped into three subgroups: <1, 1−1.65,>1.65. **(F)** Kaplan-Meier plot for TCGA samples grouped by concordance with Invasive GSCs. The dotted line indicates median survival. When considering pairwise comparisons, only the Invasive-high and Invasive-low subgroups were significantly different (p=0.0043). Further subgrouping of the TCGA samples into smaller intervals of concordance z-score yielded no benefit, preserving the Invasive-high subgroup as the only one with significantly poorer prognosis (Fig. S3D-E).

Previous work suggests that GBM tumors harbor a heterogeneous population of GSCs[14,15,28]. We therefore quantified the presence of Reactive, Constructive and Invasive cancer stem cells in our four primary GBM based on their scATAC-seq profiles. The Constructive state was dominant in every primary tumor, ranging from 9-21% of all cells captured by scATAC-seq (Fig. 4C). The Reactive and Invasive states accounted for only 0-9% of all cells, in varying proportions from one tumor to another (Fig. 4C). Collectively, our results further support the heterogeneous nature of cancer stem cells that populate primary tumors.

While various classifications of GBMs and/or their constitutive bulk and stem tumor cells have been reported, with some associating with patient survival[8,22,28–31], no molecular signature in GBM has so far been reported that can significantly stratify the poorer prognosis *IDH* wild-type patients by survival. We performed intracranial xenografts of 37 *IDH* wild-type GSC populations and classified the transplanted cells by their GSC state to perform a differential survival analysis (Fig. 4D). The overall survival times of the transplanted mice grouped by GSC state were significantly different (LogRank test p = 0.041), with the Invasive state GSCs leading to the worst prognosis. Relative to the mice injected with Reactive state GSCs, mice with Constructive state GSCs and Invasive state GSCs had hazard ratios of 1.3 (95% CI: 0.57-2.97) and 3.5 (95% CI: 1.2-10.49), respectively. Next, we investigated the prognostic value of the GSC states using the TCGA GBM cohort (*IDH* wild-type, n = 144). When classified by dominant GSC state, TCGA tumors display the same trend as the xenografts with Invasive state-dominated tumors showing the lowest survival (Fig. S3A). However, with only two tumors classified as Invasive-dominant, the difference in survival between the three patient groups was not statistically significant (p = 0.3) (Fig. S3A). We proceeded to rank the TCGA tumors solely by their concordance to Invasive GSCs and classified the patient tumors into Invasive-low (z-score < 1), Invasive-mid (z-score 1-1.65), and Invasive-high (z-score ≥ 1.65) groups (Fig. 4E). With this stratification method, median patient survival per group not only decreased with increasing Invasive GSC score, but we also identified an Invasive-high subset of tumors with significantly lower survival (p = 0.019, HR = 2.8, 95% CI: 1.3-5.81)) (Fig. 4F). Invasion assays using representative Invasive state GSCs additionally highlight the invasive properties of these populations both *in vitro* and *in vivo* (Fig. S4). These results show that cancer stem cell states defined based on the chromatin accessibility in GSCs can identify transcriptional programs associated with poor prognosis and can serve as a signature to identify high-risk patients in *IDH* wild-type GBM.

## DISCUSSION

Defining the nature of self-renewing tumor-initiating cells in primary GBM is required to identify vulnerabilities for therapeutic intervention. Quantifying their heterogeneity within tumors can guide treatment strategies and assist in predicting the course of disease progression. Here we show that chromatin accessibility assays capture a heterogeneity across self-renewing tumor-initiating cells in primary GBM that extends beyond their genetic diversity, and underlies the heterogeneity in bulk progeny[15]. This heterogeneity aligns with diversity in the three-dimensional genome organization of GSCs[32] and agrees with how the three-dimensional genome organization instructs cis-regulatory plexuses underlying gene regulation[33–37]. We further reveal a specific cancer stem state that is significantly predictive of patient survival and can be used as a signature to identify high-risk patients. Our results also highlight dependencies unique to each cancer stem state. Specifically, the Reactive GSC state-associated transcription factor SP1 and its regulatory partners are involved in cellular differentiation and growth, apoptosis, response to DNA damage, chromatin remodelling[38], stimulation of *TERT* expression in cancer stem cells[39], and increased stemness and invasion in GBM[40]. In contrast, the Constructive GSC state rely on transcription factors including OLIG2, a known GSC marker[41], AHR involved in tumor microenvironment responses and metabolic adaptation[42], NPAS3, a regulator of Notch signaling and neurogenesis[43] and ASCL1, a critical regulator of GSC differentiation and marker of sensitivity to Notch inhibition in GSCs[19,44]. Finally, the Invasive GSC state relies on FOXD1, a pluripotency regulator and determinant of tumorigenicity in GSCs regulating expression of the aldehyde dehydrogenase ALDH1A3, a functional marker for invasive GSCs[45,46]. Collectively, our results support developing combination therapy using targeting agents against each GSC state, such as Notch inhibitors[19] and small molecule inhibitors of ALDH[45], to eradicate self-renewing tumor-initiating cells with the hope to cure GBM patients.

## METHODS

### Patient samples and cell culture

All tissue samples were obtained following informed consent from patients, and all experimental procedures were performed in accordance with the Research Ethics Board at The Hospital for Sick Children (Toronto, Canada), the University of Calgary Ethics Review Board, and the Health Research Ethics Board of Alberta - Cancer Committee (HREBA). Approval to pathological data was obtained from the respective institutional review boards. Patient tumor tissue samples were dissociated in artificial cerebrospinal fluid followed by treatment with enzyme cocktail at 37°C. Patient tumor-derived GSCs were grown as adherent monolayer cultures in serum-free medium as previously described[21]. Briefly, cells were grown adherently on culture plates coated with poly-L-ornithine and laminin. Serum-free NS cell self-renewal media (NS media) consisted of Neurocult NS-A Basal media, supplemented with 2 mmol/L L-glutamine, N2 and B27 supplements, 75 μg/mL bovine serum albumin, 10 ng/mL recombinant human EGF (rhEGF), 10 ng/mL basic fibroblast growth factor (bFGF), and 2 μg/mL heparin. A subset (22/37) of the GSCs used for orthotopic xenografts were grown as non-adherent spheres prior to single-cell dissociation and injection into the mice. Briefly, serum-free medium (SFM) was used to initiate GSC cultures. Non-adherent spheres formed after 7-21 days in culture and were expanded, then cryopreserved in 10% dimethyl sulfoxide (DMSO; Sigma-Aldrich) in SFM until used in experiments.

### ATAC-seq

ATAC-seq was used to profile the accessible chromatin landscape of 27 patient tumor-derived GSCs. 50,000 cells were processed from each sample as previously described[47,48]. The resulting libraries were sequenced with 50 bp single-end reads which were mapped to hg19. Reads were filtered to remove duplicates, unmapped or poor quality (Q <30) reads, mitochondrial reads, chrY reads, and those overlapping the ENCODE blacklist. Following alignment, accessible chromatin regions/peaks were called using MACS2. Default parameters were used except for the following: --keep-dup all -B --nomodel --SPMR -q 0.05 --slocal 6250 --llocal 6250. The signal intensity was calculated as the fold enrichment of the signal per million reads in a sample over a modelled local background using the bdgcmp function in MACS2. Spectral clustering implemented in the SNFtool package[49] was run on the SNF fused similarity matrix to obtain the groups corresponding to k=2 to 12. Enrichment for genomic features was calculated using CEAS[50].

A given chromatin region was considered exclusive to one of the clusters if it was called as a peak in any of the cluster’s samples using a q-value filter of 0.05 and was not called as a peak in any of the other samples using a q-value filter of 0.2, in order to ensure stringency of exclusivity.

The ATAC-seq saturation analysis was performed by randomizing the order of samples, and successively calculating the number of additional peaks discovered with the addition of each new sample. This process was repeated 10,000 times and averaged. A self-starting non-linear regression model was then fitted to the data to estimate the level of saturation reached.

For the xenograft survival analysis, 11/37 GSCs used overlap with the cohort of 27 described above. The other 26/37 GSCs were profiled by ATAC-seq independently following the same protocol described above and assigned to a GSC state through unsupervised hierarchical clustering with the original cohort of 27 GSCs.

### Single cell ATAC-seq

The four tumors used were G4218 (primary GBM, IDH wt, Male, 64yrs), G4250 (primary GBM, IDH wt, Male, 73yrs), G4275 (primary GBM, IDH wt, Female, 52yrs), G4349 (primary GBM, IDH wt, Male, 62 yrs). Fragments of tumor were received fresh from the operating room, and blunt dissected into individual fragments of approximately 0.3-0.7 cm3. Each fragment was placed in 1 mL of freezing media (400 μL of NeuroCult NS-A Basal medium with proliferation supplement (StemCell Technologies; #05751) containing 20 μg/mL rhEGF (Peprotech, AF-100-15), 10 μg/mL bFGF (StemCell Technologies, #78003), and 2 μg/mL heparin (StemCell Technologies, #07980); 500 μL of 25% BSA (Millipore-Sigma; A9647) in DMEM, and 100 μL DMSO (Millipore-Sigma; D2650) in a 2 mL cryotube, and placed at −80 C in a CoolCell for at least 24 hours. Samples were then stored at −80C until use. Cryopreserved primary GBM samples were washed at 1000 RPM for 5 minutes in PBS to remove DMSO, and then transferred to 1.5 mL tubes. Samples were resuspended in cold ATAC resuspension buffer (10 mM Tris-HCl pH 7.4, 10 mM NaCl, 3 mM MgCl2, 0.1% NP-40, 0.1% Tween-20, 0.01% Digitonin, 1% BSA in PBS) on ice and dissociated using a wide-bore P1000 pipette tip and vortexing, followed by 10 minutes of incubation on ice. Cells were spun down at 500× g for 5 minutes at 4 C, washed in the ATAC resuspension buffer, spun down again, and resuspended in ATAC-Tween wash buffer (10 mM Tris-HCl pH 7.4, 10 mM NaCl, 3 mM MgCl2, 0.1% Tween-20, 1% BSA in PBS), then passed through a cell strainer top FACS tube (Falcon; #38030) to remove debris. Nuclei quality and quantity was evaluated using trypan blue on an Invitrogen Countess II device in duplicate, and a subset of nuclei was spun down in a fresh tube and resuspended in 10X sample dilution buffer. Nuclei were then used for single cell ATAC-seq library construction using the Chromium Single Cell ATAC Solution v1.0 kit (10X Genomics) on a Chromium controller. Completed libraries were further quality checked for fragment size and distribution using an Agilent TapeStation prior to sequencing. Single-cell ATAC-seq samples were sequenced on a NextSeq 500 (Illumina) instrument with 50 bp paired-end reads at the Centre for Health Genomics and Informatics (CHGI) at the University of Calgary.

The raw sequencing data was demultiplexed using cellranger-atac mkfastq (Cell Ranger ATAC, version 1.0.0, 10× Genomics). Single cell ATAC-seq reads were aligned to the hg19 reference genome (hg19, version 1.1.0, 10× Genomics) and quantified using cellranger-atac count function with default parameters (Cell Ranger ATAC, version 1.1.0, 10× Genomics). The resulting data were analysed using the chromVAR[51] and Signac[52] R packages (v1.4.1). The number of accessibility modules in each sample was determined using the ElbowPlot method implemented in Signac. Similarity between individual cells and GSC states was assessed using the deviation scores calculated by chromVAR within the single cell data for significantly differentially accessible sets of peaks (Fold Change Signal difference >2 and Wilcoxon test q-value <=0.05) between the states as determined by bulk ATAC-seq. Similarity between individual cells and the expression-derived cellular states was assessed using the deviation scores calculated by chromVAR within the single cell data for promoter regions of the signature genes of each of the cellular states[8]. A 2-fold cut-off was used to determine dominance of a UMAP module by an individual or group of cellular states. Similarity between individual cells and the GBM cancer stem cell signatures was assessed using the deviation scores calculated by chromVAR within the single cell data for promoter regions of the 19 transcription factors identified as markers of cancer stem cells in GBM[13].

Copy number variants in single cells were determined using CONICSmat[53] with default parameters using the gene activity matrix generated by Signac as input. We focused on chr7 gains and chr10 losses as they are hallmark chromosomal changes in GBM and found the following fractions of cells carrying these CNVs, on average across the four tumors: 76% of all cells, 88% of cells allocated to scRNAseq cellular states[8], 95% of cancer stem cells based on the 19 gene signature, 91% of GSCs based on shared accessible regions between 14/27 GSC populations, 94% of GSCs identified based on the state-specific signatures.

### DNA Methylation arrays

Bisulfite conversion of DNA for methylation profiling was performed using the EZ DNA Methylation kit (Zymo Research) on 500 ng genomic DNA from all 27 samples. Conversion efficiency was quantitatively assessed by quantitative PCR (qPCR). The Illumina Infinium MethylationEPIC BeadChips were processed as per manufacturer’s recommendations. The R package ChAMP v2.6.4[54] was used to process and analyse the data. For the copy number analysis, the raw IDAT files were uploaded to the MNP tool[25], which directly compares the copy number profile estimated from the probe intensities on the methylation array to the distribution observed across thousands of brain tumors in its database.

### RNA-seq

RNA was extracted from GSCs using the Qiagen RNeasy Plus kit. RNA sample quality was measured by Qubit (Life Technologies) for concentration and by Agilent Bioanalyzer for RNA integrity. All samples had RIN above 9. Libraries were prepared using the TruSeq Stranded mRNA kit (Illumina). Two hundred nanograms from each sample were purified for polyA tail containing mRNA molecules using poly-T oligo attached magnetic beads, then fragmented post-purification. The cleaved RNA fragments were copied into first strand cDNA using reverse transcriptase and random primers. This is followed by second strand cDNA synthesis using RNase H and DNA Polymerase I. A single “A” base was added and adapter ligated followed by purification and enrichment with PCR to create cDNA libraries. Final cDNA libraries were verified by the Agilent Bioanalyzer for size and concentration quantified by qPCR. All libraries were pooled to a final concentration of 1.8nM, clustered and sequenced on the Illumina NextSeq500 as a pair-end 75 cycle sequencing run using v2 reagents to achieve a minimum of ~40 million reads per sample. Reads were aligned to hg19 using the STAR aligner v2.4.2a [55] and transcripts were quantified using RSEM v1.2.21[56] or vst transformed using DESeq2[57].

### Motif Enrichment

Regions exclusively accessible in one of the GSC states and not the others were used as input sequences for the motif enrichment, while the full ATAC-seq catalogue served as the background set when running HOMER v4.7 to detect enrichments of transcription factor binding motifs. Enriched motifs were then grouped into families based on similarities in DNA-binding domains using the CIS-BP database[58]. Each family was assigned the fold-enrichment value of the most enriched motif within the family.

The transcription factors whose motifs were found enriched in Reactive-exclusive accessible regions, were run together through GSEA[23], and the gene set corresponding to genes potentially regulated by SP1 was identified as significantly enriched (GSEA gene set GGGCGGR_SP1_Q6). The expression levels of key transcription factors in tumor and normal samples were analysed and displayed using GEPIA[27].

### Gene essentiality screen

Illumina sequencing reads from genome-wide TKOv1 CRISPR screens in patient-derived GSCs[26] were mapped using MAGECK[59] and analysed using the BAGEL algorithm with version 2 reference core essential genes/non-essential genes[60,61]. Resultant raw Bayes Factor (BF) statistics were used to determine essentiality of transcription factor genes using a minimum BF of 3 and a 5% FDR cut-off. For visualisation purposes only, the essentiality scores were scaled and the individual GSC essentiality thresholds subtracted from each score to obtain a common threshold at 0 across GSCs.

### Orthotopic xenografts

All animal procedures were performed according to and approved by the Animal Care Committee of the Hospital for Sick Children or the University of Calgary. All attempts are made to minimize the handling time during surgery and treatment so as not to unduly stress the animals. Animals are observed daily after surgery to ensure there are no unexpected complications. For intracranial xenografts, 100,000 GSC cells were stereotactically injected into the frontal cortex of 6-8 weeks old female NOD/SCID or C17/SCID mice. Mice were monitored and euthanized once neurological symptoms were observed or at the experimental endpoint of 12 months.

### Invasion assay

Hydrogels were synthesized as previously described[62], with the following modifications: 1% w/v hyaluronan-methyl furan, 2.3mM MMP cleavable crosslinker, and 400μM fibronectin-derived peptide with sequence Mal-SKAGPHSRNGRGDSPG. Cells were plated on hydrogels at a density of 3500 cells/hydrogel and allowed to adhere for 24 h. 48 h after seeding, fresh media was added to each well. Cells were fixed with 4% PFA 4-5 days after seeding. Cells were counterstained with Hoechst (1/500) to label nuclei, Alexa FluorTM 488 Phalloidin (1/40) to label F-actin, and 15 μm FluosphereTM red beads (1/15) to label the surface of the hydrogel. Hydrogels were imaged using confocal microscopy at 10x magnification, taking images every 20 μm on the z-axis. Imaris Bitplane 8.3.1 software was used to prepare images and to analyze the positions of each cell and surface bead label. A custom Matlab script was used to calculate the percent invasion, defined as the percentage of total cells located below a 75 μm threshold from the surface of the hydrogel. Statistics were calculated using Graphpad Prism 7.04 software. Plots are shown as mean with standard deviation, and statistics shown are from one-way ANOVA with multiple comparisons with Dunnet’s test correction, comparing all patient cell lines (dark grey) to the hf6562 healthy control. Data were displayed with p values represented as * p ≤ 0.05, ** p ≤ 0.01, *** p ≤ 0.001, **** p ≤ 0.0001.

### Immunohistochemistry

Tissue samples were formalin fixed and paraffin embedded. Serial sections deparaffinized, rehydrated through an alcohol gradient to water and antigen retrieval in citrate buffer pH 6.0 was used for the human nucleolin antibody at 5.0 g/mL (ab13541) (Abcam, Canbridge, MA). Endogenous peroxide activity and nonspecific binding was blocked with 3%(v/v) peroxide and 2% (v/v) normal horse serum. Primary antibody and anti-mouse ImmPRESS-HRP secondary antibody were incubated for 1 hr and visualized using DAB (3,3’-diaminobenzidine) (Vectorlabs, Burlingame, CA). Normal horse serum or monoclonal IgM was used in control sections.

### Survival analysis

Survival analysis on xenografts and TCGA data was performed using R packages survival[63] and survminer[64]. The LogRank test was used in every analysis. See ATAC-seq section for details on how each GSC used in the orthotopic xenografts was assigned to a GSC state. TCGA samples were assigned to individual GSC states in the following way. 1) Using the unsupervised clustering of RNA-seq data presented in Fig. 3B, the 23/27 GSCs that displayed matched GSC state assignments by RNA-seq and ATAC-seq were used in this analysis. 2) Genes preferentially enriched in each GSC state were determined using DEseq2[57] (q ≤ 0.05 and fold-change ≥ 2). 3) The mean log2(FPKM+1) value for each of these genes over all GSCs in each state was calculated to obtain a single representative value for each gene in each of the three GSC states. 4) the concordance index was then calculated between each TCGA sample and each GSC state and individual TCGA samples were assigned to the GSC state with the highest score. Similarly, to assign TCGA samples to the three Invasive groups (Invasive-low, -mid, and -high), the concordance to Invasive GSCs as calculated above was used. The z-score for each sample was then used to classify each TCGA sample into the three subgroups of Invasive-low (Invasive z-score < 1), Invasive-mid (Invasive z-score 1-1.65), and Invasive-high (Invasive z-score ≥ 1.65). When changing the Invasive z-score thresholds for grouping the TCGA samples, the most Invasive-high subgroup remains associated with the lowest survival (Fig. S3B-C).

### Data and materials availability

The GSCs are available upon reasonable request from PBD and SW. The GSC ATAC-seq and DNA methylation data have been deposited at GEO. The scATAC-seq data has been deposited at GEO. RNA-seq data are available at EGA.

### Code availability

All data analysis was performed using established methods implemented in published software or R packages. Software and package versions and parameters are detailed in the Methods section. All scripts used for the analysis are available upon request.

## ACKNOWLEDGMENTS

We would like to thank the Princess Margaret Genomics Centre for their assistance in this study and 10x Genomics, especially Adam Jerauld, Ariel Royall and John Chevillet for training and support. We also thank Aude Gerbaud and Stacey Krunholtz for their help with figure formatting and David Vetrie for his thoughtful input on the manuscript. This work was supported by the Princess Margaret Cancer Foundation, the Canadian Epigenetics, Environment, and Health Research Consortium (CEEHRC) initiative from the Canadian Institute for Health Research (CIHR) (TGH-158221 to ML, SA and PBD) and SU2C Canada Cancer Stem Cell Dream Team Research Funding (SU2C-AACR-DT-19-15) provided by the Government of Canada through Genome Canada and the Canadian Institute of Health Research, with supplemental support from the Ontario Institute for Cancer Research, through funding provided by the Government of Ontario. Stand Up To Cancer Canada is a Canadian Registered Charity (Reg. # 80550 6730 RR0001). Research Funding is administered by the American Association for Cancer Research International - Canada, the Scientific Partner of SU2C Canada. This study was also conducted with the support of the Ontario Institute for Cancer Research through funding provided by the Government of Ontario for the Brain Cancer Translational Research Initiative. PG was supported by a CIHR Fellowship (MFE 338954). Funding to HAL and SW was from CIHR. MG holds a Canada Research Chair in Brain Cancer Epigenomics and is supported by CIHR, NSERC and the Alliance for Cancer Gene Therapy. PBD is also supported by CIHR, OICR, the Terry Fox Research Institute, and the Hospital for Sick Children Foundation, Jessica’s Footprint, Hopeful Minds, and the Bresler Family. PBD holds a Garron Chair in Childhood Cancer Research at the Hospital for Sick Children. SA is supported by the CIHR, the Terry Fox Institute and the Canadian Cancer Society. ML holds an Investigator Award from the Ontario Institute for Cancer Research and a Canadian Institutes of Health Research (CIHR) New Investigator Award.

## AUTHOR CONTRIBUTIONS

ML, PG, and PBD conceptualized and designed the study assisted by FJC. PG conducted the genomics experiments, designed and/or implemented most of the computational and statistical approaches, and made the figures. MMK and RH performed all the tissue culture, under the supervision of PBD, HAL and SW. AN, KE, and DS generated the scATAC-seq data from samples provided by JAC, under the supervision of MG. BN performed the alignment of the scATAC-seq data. NR, XH and RH performed the xenografts, under the supervision of PBD, HAL, and SW. CA generated some of the ATAC-seq data used in the xenograft classification. FMGC ran the spectral clustering, under the supervision of MDT. GM, NR and SA contributed the essentiality screen data. HW conducted the endpoint xenograft staining. LJS performed the invasion assay under the supervision of MSS. SAMT contributed to the computational analysis design, under supervision of BHK and ML. The manuscript was written by PG, PBD and ML with input from all other authors.

## COMPETING INTERESTS

The authors declare no competing interests.

## SUPPLEMENTARY DATA

**Fig. S1. (A-D)** UMAPs showing cells confirmed to be GBM tumor cells, with gains of chr7 or losses of chr10. **(E-G)** Overlaid ATAC signal tracks for the 27 GSCs at the 92 most differentially accessible regions between GSC states as determined by pairwise wilcoxon test and a median signal fold change equal or greater than 2. (E) Most accessible in Reactive state GSCs. (F) Most accessible in Constructive state GSCs. (G) Most accessible in Invasive state GSCs.

**Fig. S2. (A)** The log2(fold enrichment) for all motif families were ordered and plotted. A cut-off threshold to select families for follow-up was selected based on the steep inflection of the curve above 0.5. **(B)** Schematic of drop-out essentiality screen using GSCs stably expressing Cas9 and gRNA libraries. **(C-E)** The expression values determined by RNA-seq for all transcription factors whose motif is enriched in each GSC state and exclusively essential in at least one GSC of each state is plotted. Bold colors highlight factors essential in all tested GSCs in that state. In the case of the Reactive state, blue boxplots correspond to SP1 and members of its regulatory network. **(F)** Expression levels of key transcription factors in tumor and normal samples, analysed and displayed using GEPIA[27]. **(G)** Gene enrichment analysis of subtype-specific essential genes (Essentiality), upregulated and differentially accessible genes (ATAC and RNA combined), and putative targets of key transcription factors.

**Fig. S3. (A)** Kaplan-Meier plot for TCGA samples grouped by GSC state. The dotted lines indicate median survival. **(B)** TCGA samples ordered by increasing concordance with Invasive GSCs and grouped into 8 subgroups. **(C)** Kaplan-Meier plot for TCGA samples grouped by concordance with Invasive GSCs. The dotted line indicates median survival.

**Fig. S4. (A)** Left: Top and side views of confocal z-stack images of Invasive GSCs G705, G837 and healthy control hf6562 cultured on hyaluronan-based hydrogels. Right: percent invasion, defined as the percentage of total cells located below a 75 μm threshold from the surface of the hydrogel. Plots are shown as mean with standard deviation, and statistics shown are from one-way ANOVA with multiple comparisons with Dunnet’s test correction, comparing all patient cell lines (dark grey) to the hf6562 healthy control. Data were displayed with p values represented as * p ≤ 0.05, ** p ≤ 0.01, *** p ≤ 0.001, **** p ≤ 0.0001. **(B)** Human-specific staining of mouse brains injected with GSCs from G411 (upper panel) and G837 (lower panel) with higher magnification (black inset) at the mass boundaries.

