## Supplementary figures and images for "Single-cell chromatin accessibility in glioblastoma delineates cancer stem cell heterogeneity predictive of survival"

### Supplementary material

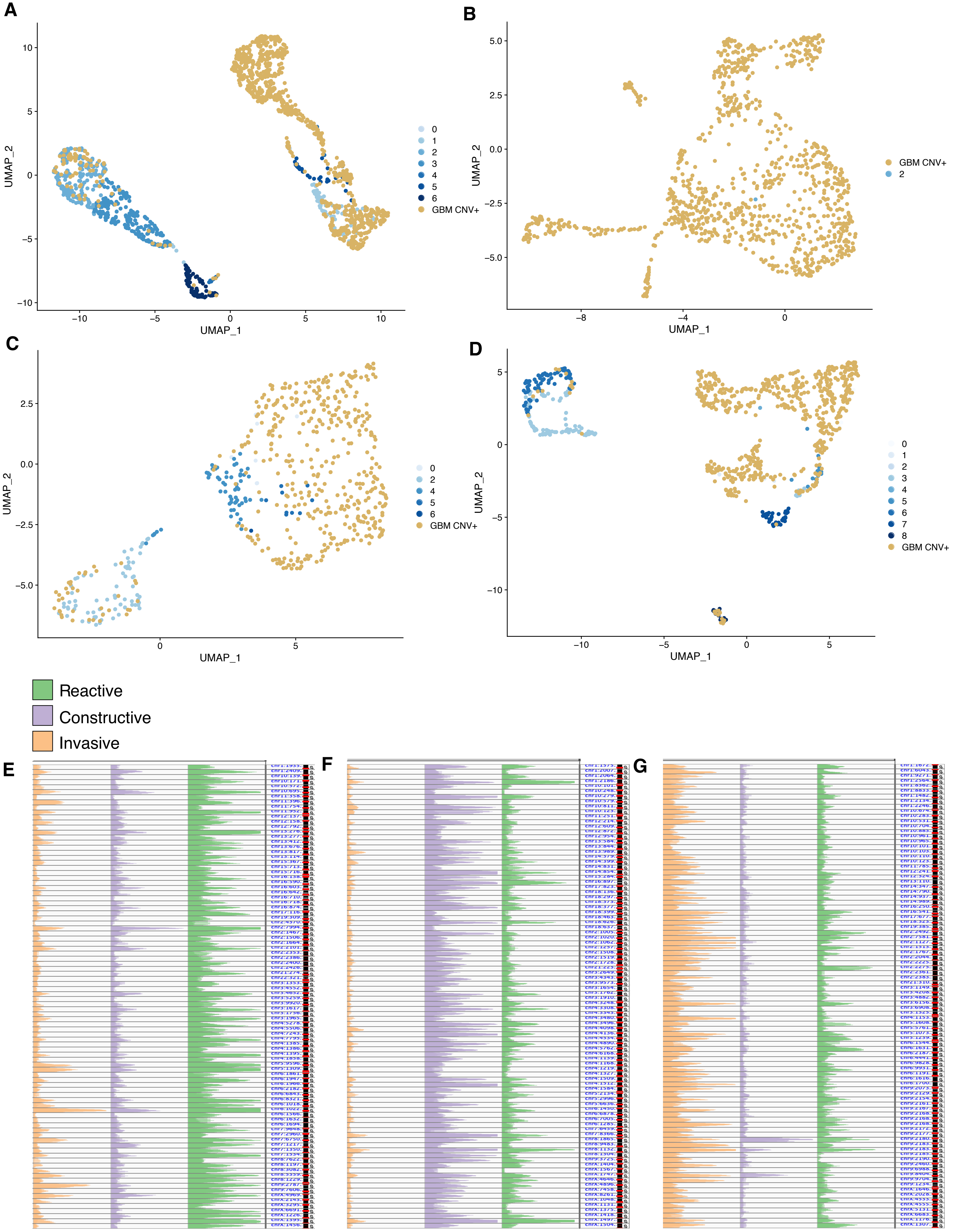

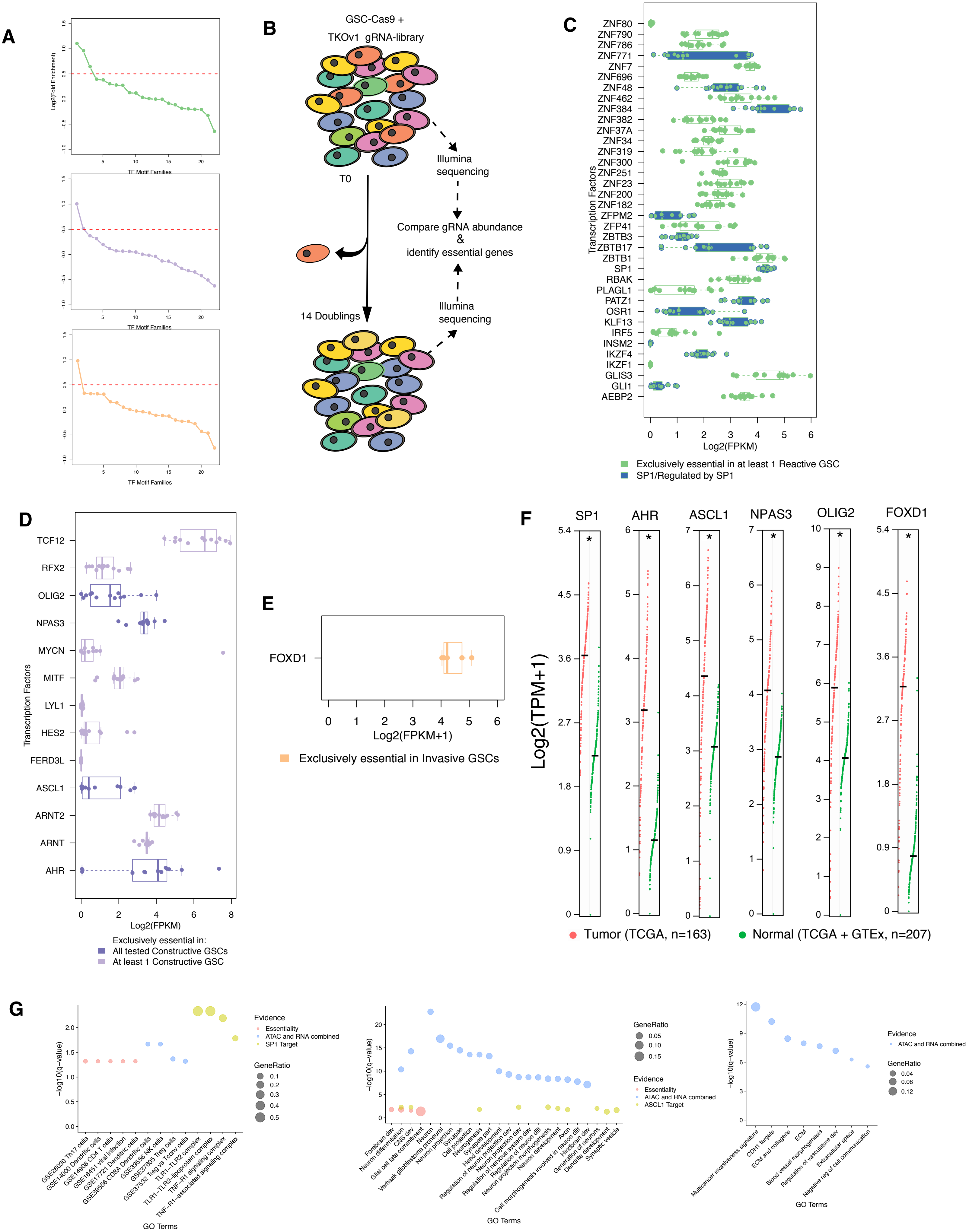

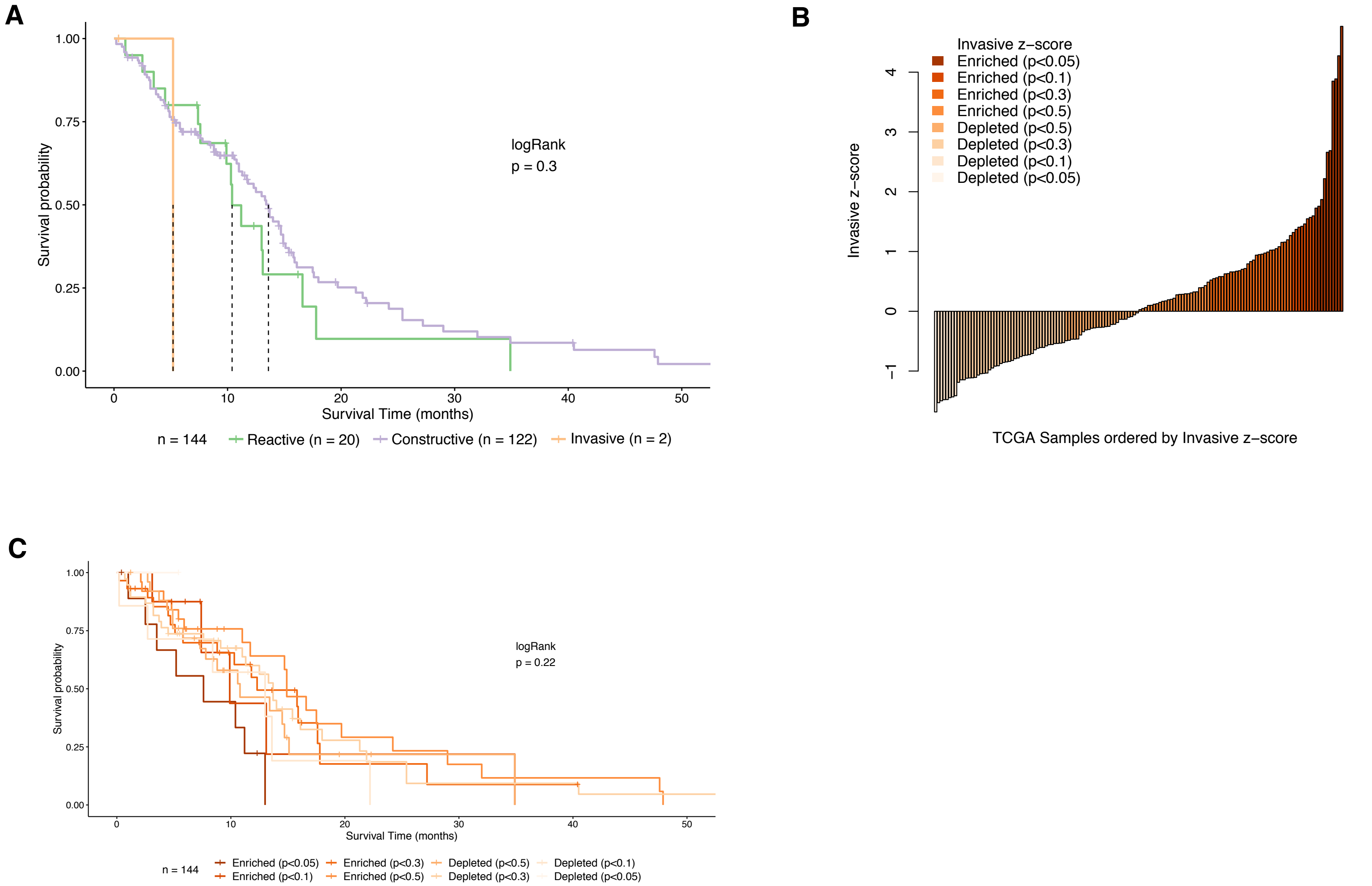

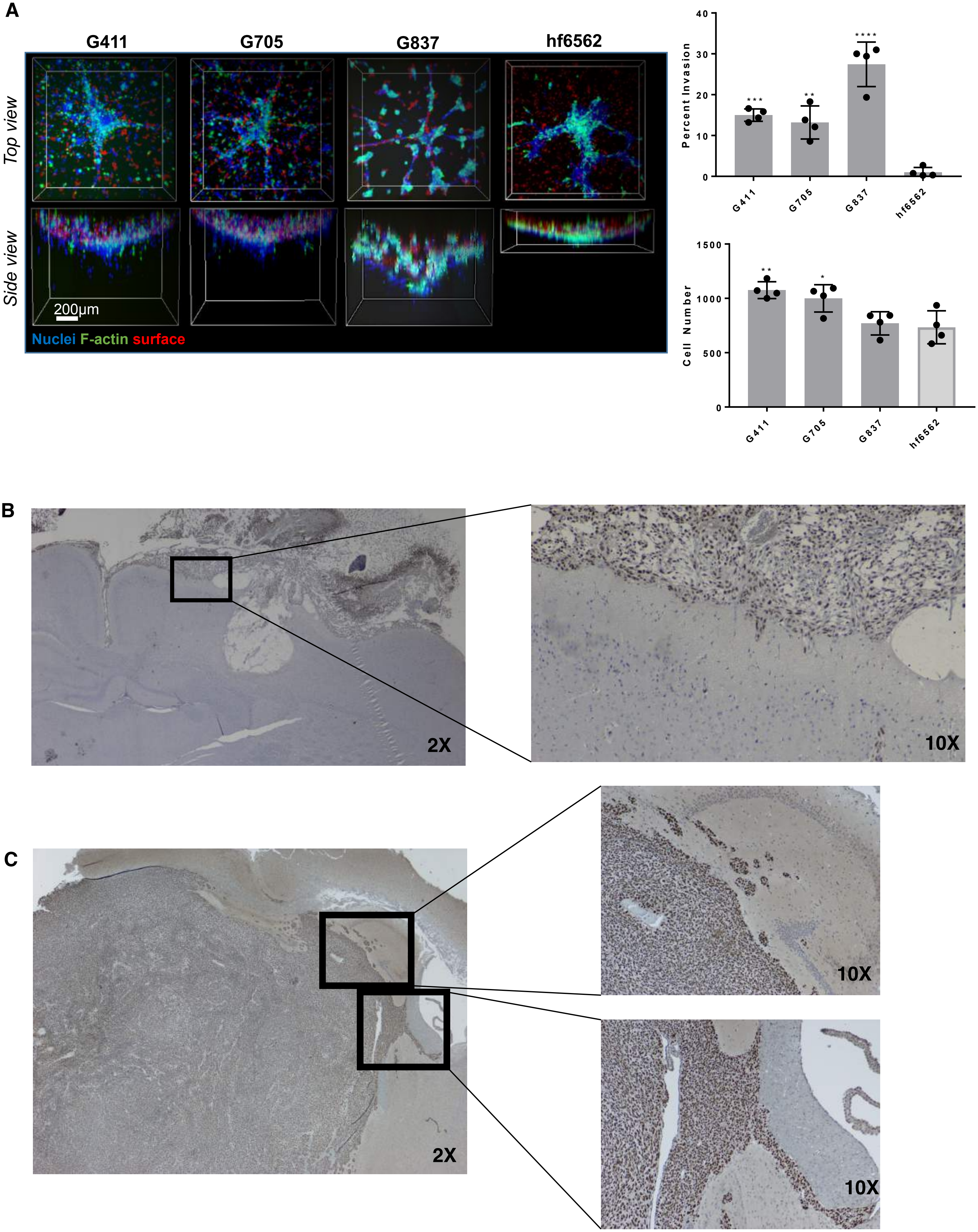
